# A phenome-wide examination of neural and cognitive function

**DOI:** 10.1101/059733

**Authors:** RA Poldrack, E Congdon, W Triplett, KJ Gorgolewski, KH Karlsgodt, JA Mumford, FW Sabb, NB Freimer, ED London, TD Cannon, RM Bilder

**Author notes:** Corresponding Author: R Poldrack.

## Abstract

This data descriptor outlines a shared neuroimaging dataset from the UCLA Consortium for Neuropsychiatric Phenomics, which focused on understanding the dimensional structure of memory and cognitive control (response inhibition) functions in both healthy individuals (138 subjects) and individuals with neuropsychiatric disorders including schizophrenia (58 subjects), bipolar disorder (49 subjects), and attention deficit/hyperactivity disorder (45 subjects). The dataset includes an extensive set of task-based fMRI assessments, resting fMRI, structural MRI, and high angular resolution diffusion MRI. The dataset is shared through the OpenfMRI project, and is formatted according to the Brain Imaging Data Structure (BIDS) standard.

## Background and summary

The Consortium for Neuropsychiatric Phenomics (CNP) was established on the principle that discovery of genetic mechanisms for complex mental disorders will ultimately demand integration of research on large numbers of *phenotypes* spanning multiple biological and behavioral scales, from genome to syndrome, and will require both novel informatics and data analytic strategies. The CNP focused on two phenotype domains – memory/working memory and response inhibition – because these neurocognitive domains have been examined across multiple levels of analysis and are relevant to multiple neuropsychiatric syndromes. To examine these domains the CNP collected interviews and rating scales, self-report measures, neurocognitive exams using both paper-pencil and computerized tests, and neuroimaging data comprising structural MRI (sMRI), high angular resolution diffusion imaging (HARDI), and functional MRI (fMRI) both at rest (rsfMRI) and during five different cognitive activation paradigms. The CNP was committed to the perspective that most phenotypes should be considered dimensional until evidence of categorical structure was clear, and for this reason focused most on assessing phenotypes across a broad range of healthy people rather than specific groups representing individuals with neuropsychiatric syndromes. The CNP sample additionally included smaller groups of people who had diagnoses of schizophrenia, bipolar disorder, and attention-deficit/hyperactivity disorder (following DSM-IV), to determine where these individuals’ scores would fall with respect to the distributions observed in healthy individuals. The CNP was one of 9 Interdisciplinary Research Consortia supported by the NIH Roadmap Initiative from 2007-2012. The CNP comprised 8 linked awards, including a Coordinating Center (UL1DE019580, UL1RR024911), five linked R01 awards (RL1MH083268, RL1MH083269, RL1DA024853, RL1MH083270, and RL1LM009833) and two center grant awards (PL1MH083271 and PL1NS062410). To date the neuroimaging data from the CNP study have been used in 6 publications ^1–6^, and another 53 publications are linked to CNP awards.

This Data Descriptor provides a description of the neuroimaging data and related data from the CNP, which have been shared via the OpenFMRI project (http://openfmri.org)^78^. The shared data are described according to the Brain Imaging Data Structure (BIDS: http://bids.neuroimaging.io) (Gorgolewski et al, in press) which is a data organization format designed to facilitate the transfer, description, and storage of neuroimaging experiment data. BIDS provides a disciplined way to arrange the different data types that comprise a neuroimaging experiment.

## Methods

### Participants

Healthy adults, ages 21-50, were recruited by community advertisements from the Los Angeles area; adult ADHD, Bipolar and Schizophrenia participants were recruited using a patient-oriented strategy involving outreach to local clinics and online portals (separate from the methods used to recruit healthy volunteers). All candidates were screened by telephone and then in person. To be included individuals had to identify themselves in either one of the NIH ethnic and racial categories: White, not Hispanic or Latino; or Hispanic or Latino, of any racial group. Additionally they satisfied the following inclusion-exclusion criteria: primary language (as determined by verbal fluency tests in both languages) either English or Spanish; completed at least 8 years of formal education; no significant medical illness by self report; adequately cooperative to complete assessments; and visual acuity 20/60 or better. Urinalysis was used to screen for drugs of abuse (cannabinoids, amphetamine, opioids, cocaine, benzodiazepines), and participants were excluded if results were positive. This healthy behavioral sample comprised 1101 individuals.

A subset of the healthy sample and patient sample took part in two separate fMRI sessions, which each included one-hour of behavioral testing and a one-hour scan on the same day. Eligible English-speaking participants between the ages of 21 and 40 were recruited from the parent study if they successfully completed all previous testing sessions, and they did not meet the following additional exclusion criteria: left-handedness, pregnancy, history of head injury with loss of consciousness or cognitive sequelae, or other contraindications to scanning (e.g., claustrophobia, metal in body).

After receiving a verbal explanation of the study, participants gave written informed consent following procedures approved by the Institutional Review Boards at UCLA and the Los Angeles County Department of Mental Health. All subjects underwent a semi-structured assessment with the Structured Clinical Interview for the Diagnostic and Statistical Manual of Mental Disorders, Fourth Edition (DSM-IV) (SCID-I; (First MB, 2004)), supplemented for ADHD diagnoses with the Adult ADHD Interview (a structured interview form derived from the Kiddie Schedule for Affective Disorders and Schizophrenia, Present and Lifetime Version (KSADS-PL) (Kaufman et al., 1997)), in order to enable a more detailed characterization of lifetime history of ADHD in adults.

The final sample included imaging data for healthy individuals from the community (138 subjects), as well as samples of individuals diagnosed with Schizophrenia (58), Bipolar Disorder (49), and ADHD (45).

### Behavioral assessment

Enrolled participants completed extensive neuropsychological testing; these data will be shared separately at a later date via the NIH Data Archive. The list of tests performed during the behavioral session is presented in Table S1. Participants were debriefed and compensated for their time at the end of the final testing session.

### Neuroimaging assessment

*Imaging acquisition*. MRI data were acquired on one of two 3T Siemens Trio scanners, located at the Ahmanson-Lovelace Brain Mapping Center (Siemens version syngo MR B15) and the Staglin Center for Cognitive Neuroscience (Siemens version syngo MR B17) at UCLA. Functional MRI data were collected using a T2*-weighted echoplanar imaging (EPI) sequence with the following parameters: slice thickness = 4 mm, 34 slices, TR = 2 s, TE = 30 ms, flip angle = 90°, matrix 64 × 64, FOV = 192 mm, oblique slice orientation. Additionally, a T2-weighted matched-bandwidth high-resolution anatomical scan (with the same slice prescription as the fMRI scan) and MPRAGE were collected. The parameters for the high-resolution scan were: 4mm slices, TR/TE=5000/34 ms, 4 averages, matrix = 128×128, 90 degree flip angle. The parameters for MPRAGE were the following: TR = 1.9 s, TE = 2.26 ms, FOV = 250 mm, matrix = 256 × 256, sagittal plane, slice thickness = 1 mm, 176 slices. Diffusion weighted imaging (DWI) data were collected using an echo-planar sequence with parameters: 64 directions, 2mm slices, TR/TE=9000/93 ms, 1 average, 96=96 matrix, 90 degree flip angle, axial slices, b=1000 s/mm^2^.

Subjects participated in two scanning sessions ("A" and "B") in a counterbalanced fashion. Session A included: localizer, MPRAGE, DWI, Matched Bandwidth Hires, Balloon Analog Risk Task, and Paired-associate Memory Task. Session B included: localizer, Matched Bandwidth Hires, Resting State, Breath Hold Task, Stop Signal Task, Spatial Capacity working memory task (SCAP), and Task Switching Task.

*Resting fMRI*. The resting fMRI scan lasted 304 seconds. Participants were asked to remain relaxed and keep their eyes open; they were not presented any stimuli or asked to respond during the scan.

*Task fMRI*. For each task described below, all participants received brief training on each task immediately before scanning. Each participant viewed the task stimuli through MRI-compatible goggles, responded with his or her right hand on an MR-compatible button box in the scanner, and performed one run of each task while in the scanner. The presentation and timing of all stimuli and response events were achieved using Matlab (Mathworks) and the Psychtoolbox (www.psychtoolbox.org) on an Apple Powerbook. Code for all tasks is available at https://poldracklab.stanford.edu/software.

*Balloon Analog Risk Task*. In this task (Lejuez et al., 2002), participants were allowed to pump a series of virtual green (experimental) and white (control) balloons. On each trial, participants chose to pump the balloon or cash out and collect their accumulated earnings for that round. For experimental balloons, after a trial in which the participant successfully pumped the balloon (meaning it did not result in an explosion), an image of a larger balloon was presented, the participant earned 5 points, and was able to pump again or cash out. After a trial in which the participant chose to cash out, the participant’s accumulated earnings for that round were displayed and the task moved onto the next round. On an explosion trial (necessarily following a pump trial), an exploded balloon was presented, the participant received no points for that round, and the task moved onto the next round. In this version of the BART, balloons exploded randomly on a number drawn from a uniform distribution over numbers of pumps, with 12 maximum pumps possible before an explosion or end of a round. Thus, participants experienced the probability as non-stationary, as the likelihood of a loss event increased with each trial in a round and as no information was provided to subjects about the probability of explosion. Participants also responded to control balloons, which increased in size on successive trials, but which neither resulted in points nor exploded. For both balloons (green and white), the balloon would disappear from the screen once the participant responded, and each balloon trial was separated by a jittered delay. An outcome trial (following a cash-out trial or an explosion) was displayed for a fixed duration of 2 s. Each trial was separated by a blank screen that was presented for a variable duration (1-2 s, average 1.5 s); each round was separated by a blank screen that was presented for variable duration (1-12 s, average 4 s).

The task performed in the scanner was self-paced, but the task was programmed such that participants saw approximately 30 virtual balloons, with an approximate run time of 9 minutes. Each successful pump was worth 5 points, but participants did not collect their earnings at the end of the scan.

*Paired-associate Memory Task*. In this task, two scans were performed to assess declarative memory encoding and retrieval, respectively. The first scan consisted of a block of 64 encoding trials. Forty of the encoding trials were “memory” trials and 24 were “control” trials. During memory trials, first, two words appeared for 1 s, one on each side of the screen. Then, line drawings of two objects that matched the words appeared above the words and they were presented together for 3 more seconds. One of the objects was drawn in black and white, and one object drawn using a single color (e.g. orange). For control trials, pairs of scrambled stimuli, one black and white and one colored, appeared for 2 s. The subject indicated by button press which side the colored object was on. Subjects were instructed to remember the objects and the relationship between the objects. The trial types were intermixed, and the ITI between trials was jittered. The encoding run, in total, was 8.07 minutes long.

During the retrieval scan there were 104 total trials: 24 control trials, 40 correct trials, and 40 incorrect trials. The retrieval task required the subjects to look at a pair of objects and rate their confidence in their memory of the pairing. There were 4 possible response options ranging from "Sure correct" to "Sure incorrect", allowing the responses to be analyzed as a spectrum or binarized into yes/no type responses. During control trials, on one side of the screen was one of the four retrieval confidence response options "sure correct", "maybe correct", "maybe incorrect", or "sure incorrect". On the other side of the screen was "xxxx". Subjects were asked to press the button (1-4) that corresponded to the response option displayed (for example, a response of 1 if “sure correct” appeared). In the 40 correct trials, items were shown paired as they had been at encoding. During the 40 incorrect trials items were shown paired differently than they were at encoding; some objects are the same, but were just paired incorrectly. The retrieval scan was 8.93 minutes long.

*Spatial Working Memory Task*. During the spatial delayed response task (or, spatial capacity task - SCAP), subjects were shown a target array of 1, 3, 5 or 7 yellow circles positioned pseudorandomly around a central fixation cross. After a delay, subjects were shown a single green circle and were required to indicate whether that circle was in the same position as one of the target circles had been. A relatively long stimulus presentation time of two seconds was used to allow subjects to fully encode the target array, minimizing a potential encoding bias on the basis of set size interaction. Likewise, decision or selection requirements were kept constant across set sizes to reduce possible effects of set size on response processes. In addition to load, delay period was manipulated, with delays of 1.5, 3 or 4.5 seconds. Trial events included a 2-sec target-array presentation, a 1.5, 3 or 4.5 sec delay period, and a 3-sec fixed response interval. A central fixation was visible throughout each of the 48 trials (12 per memory set size, with 4 at each delay length for each memory set). Half the trials were true-positive, and half were true-negative. Before starting the in-scanner task, subjects underwent a supervised instruction and training period outside of the scanner, and once in the scanner were again reminded of the instructions. (Glahn, 2003; Cannon, 2005)

*Stop Signal Task*. Participants were instructed to respond quickly when a “go” stimulus was presented on the computer screen, except on the subset of trials where the “go” stimulus was paired with a “stop” signal (Figure 1). Specifically, participants were shown a series of go stimuli (left-and right-wards pointing arrows), to which participants were told to respond with left and right button presses, respectively (Go trials). On a subset of trials (25%), a stop-signal (a 500 Hz tone presented through headphones) was presented a short delay after the go stimulus appeared and lasted for 250 ms (Stop trials). Participants were instructed to respond as quickly and accurately as possible on all trials, but to withhold their response on Stop trials (on trials with the tone). They also were instructed that stopping and going were equally important.

On Stop trials, the delay of the onset of the stop-signal, or stop-signal delay (SSD), was varied, such that it was increased after the participant successfully inhibited in response to a stop-signal (making the next stop trial more difficult), and decreased after the participant failed to inhibit in response to a stop-signal (making the next stop trial less difficult). Each SSD increase or decrease was in 50-ms intervals. The SSD values were drawn from two interleaved staircases per block, resulting in 16 trials from each staircase for a total of 32 Stop trials per block. On the testing day, participants completed two experimental runs (one run outside of the scanner and one while inside of the scanner). In the first task run completed outside of the scanner, SSD values started at 250 and 350 ms for staircase 1 and 2, respectively. At the end of the behavioral run, the last SSD time from each staircase was then carried over to be the initial SSD for the scan run. This one-up/one-down tracking procedure ensured that subjects successfully inhibited on approximately 50% of inhibition trials. Also as a result, difficulty level is individualized across subjects and both behavioral performance and numbers of successful stop trials are equated across subjects.

Each experiment run contained 128 trials, 96 of which were Go trials and 32 of which were Stop trials, each presented randomly. All trials were preceded by a 500 ms fixation cross in the center of the screen, then each trial began with the appearance of an arrow and ended after 1000 ms, followed by the null period. Jittered null events separated every trial (with a blank screen), with the duration of null events sampled from an exponential distribution (null events ranged from 0.5 to 4 s, with a mean of 1 s).

*Task-switching Task*. In this task, participants were presented with a series of one of four possible stimuli and asked to respond to the stimulus based on the task cue presented prior to, and above, the image. The four stimuli included a red triangle, red circle, green triangle, and green circle. Participants switched between responding to the image’s color (i.e., red or green) or the shape (i.e., triangle or circle). Cues presented included either “SHAPE” or “S” on trials where participants were expected to respond to the shape of the stimulus; cues presented included either “COLOR” or “C” on trials where participants were expected to respond to the color of the stimulus. On 33% of trials the instructions switched, such that participants were instructed to switch from responding from shape to color, or vice versa. On 67% of trials, the instructions remained the same but the cues changed. This task is designed to measure the changes in reaction time between trials requiring versus not requiring a switch in responding. Participants completed a total of 96 trials, for a total run time of 6 min 52 s.

*Breath Holding Task*. In this task, participants were asked to alternate between holding their breath and breathing regularly while resting. Participants were presented with bars on the screen in order to pace breath holding. When the bar was green, they were asked to breathe regularly; when the bar was yellow, they were asked to prepare to hold their breath; and when the bar was red, they were asked to hold their breath for 13.5 seconds. This cycle was repeated eight times, for a total run time of 2 minutes and 46 seconds. Participants performed the BHT in the scanner while wearing a respiratory belt in order to measure breathing. The purpose of this task was to assess the contribution of respiratory rhythms to changes in the BOLD signal.

*Physiological recording*. During the breath holding and resting fMRI scans, physiological data were collected using a BIOPAC MP150 with Pulse Oximeter and Respiration modules. Data were sampled at 1000 Hz and recorded using BIOPAC Acknowledge software.

## Data Records

The data set is hosted on OpenfMRI under the accession number ds000030. The files are organized in the Brain Imaging Data Structure (BIDS) version 1.0.0rc3 format, and provided in multiple gzip-compressed tar archives ^8^. BIDS provides a consistent naming convention and folder structure that is amenable to automated processing.

At the top-level, machine-readable text files in javascript object notation (JSON) are provided describing each fMRI task. A demographics file (participants.tsv; tab-delimited format) contains the unique participant number, age group, diagnosis, and an indication of which scan data are present for each subject.

The neuroimaging data collected for each participant are organized in a *sub-<subject_id>* data folder. The data for each imaging series are further organized into subfolders reflecting their intended use or purpose:

- **func**: BOLD contrast fMRI image data (both task-based and resting state), fMRI task event files, physiological monitoring logs;
- **anat:** high resolution T1-weighted image data used for structural analysis, cortical parcellation, and co-registration to standard templates;
- **dwi:** Diffusion-weighted image data used for experimental white matter characterization methods such as Diffusion Tensor Imaging;
- **beh:** Stop Signal task training run collected outside of the MRI scanner.

Each of these folders contain the following data files that collectively represent a participant’s study visit:

- **func/sub-<subject id>_task-<taskname>_bold.nii.gz**: fMRI time series image data collected from the MRI in NIfTI format and compressed with gzip;
- **func/sub<subject id>_task-<taskname>_events.tsv**: Tab-separated text file containing the timing of the events presented during the fMRI task and the responses given by the participant;
- **func/sub-<subject id>_task-<taskname>_bold.json**: a JSON-structured file containing the MRI scanner parameters used to collect the image data;
- **func/sub-<subject id>_task<taskname>_physio.tsv.gz**: A gzip-compressed tab-separated text file containing the output from physiological monitoring that occurred during the fMRI resting state and breath-holding fMRI experiments;
- **dwi/sub-<subject id>_dwi.bval**: Diffusion weighting (s/mm^2^) applied to each volume in the diffusion weighted image series;
- **dwi/sub-<subject id>_dwi.bvec**: Diffusion weighting gradient orientations applied to each volume in the diffusion weighted image series;
- **dwi/sub-<subject id>_dwi.nii.gz**: Diffusion image data in NIfTI format compressed with gzip;
- **dwi/sub-<subject id>_dwi.json**: Diffusion weighted imaging scan parameters and metadata in a JSON text file;
- **anat/sub-<subject id>_T1w.nii.gz**: High-resolution (1×1×1 mm resolution) Tl-weighted image data for structural/anatomical use;
- **anat/sub-<subject id>_T1w.json**: MR scanning parameters and metadata in a JSON text file.

The label **<taskname>** refers to data from one of the tasks performed during the visit:

- **bart**: Balloon analog risk task
- **stopsignal**: Stop signal task
- **pamenc**: Paired memory encoding task
- **pamret**: Paired memory retrieval task
- **scap**: Spatial working memory task
- **taskswitch**: Task switching task
- **bht**: Breath-holding task
- **rest**: Resting state (eyes open)

## Technical Validation

Imaging files were converted from primary DICOM data to Neuroimaging Informatics Technology Initiative (NIfTI) version 1.1 format using the dcm2niix (https://github.com/neurolabusc/dcm2niix) program. dcm2niix extracts the image pixel data and pertinent metadata parameters from the DICOM files to populate the NIfTI file header. For diffusion weighted images, dcm2niix also extracts the diffusion gradient strengths and orientations to separate files. The dcm2niix program output was saved for each conversion and inspected for errors. Additional metadata were extracted from the DICOM files using the gdcm library (http://gdcm.sourceforge.net/wiki/index.php/Main_Page) and converted to javascript object notation (JSON) format to accompany the respective image files.

All relevant metadata were combined and plotted over the timeframe of the study to reveal instances where individual scans were performed with MRI parameter settings that deviated from the standard protocol and to highlight scans with non-standard dimensions. There were very few deviations, and most were minor in nature that did not warrant data exclusion; however the table of parameters and plots are included with the data set as reference material.

After processing the source data, BOLD contrast (fMRI) and Tl-weighted anatomical imaging data were processed by the MRI Quality Control protocol (MRIQC https://github.com/poldracklab/mriqc). MRIQC computes several quality control metrics available in the published literature.

### Anatomical T1w Scans

**Figure 1:**
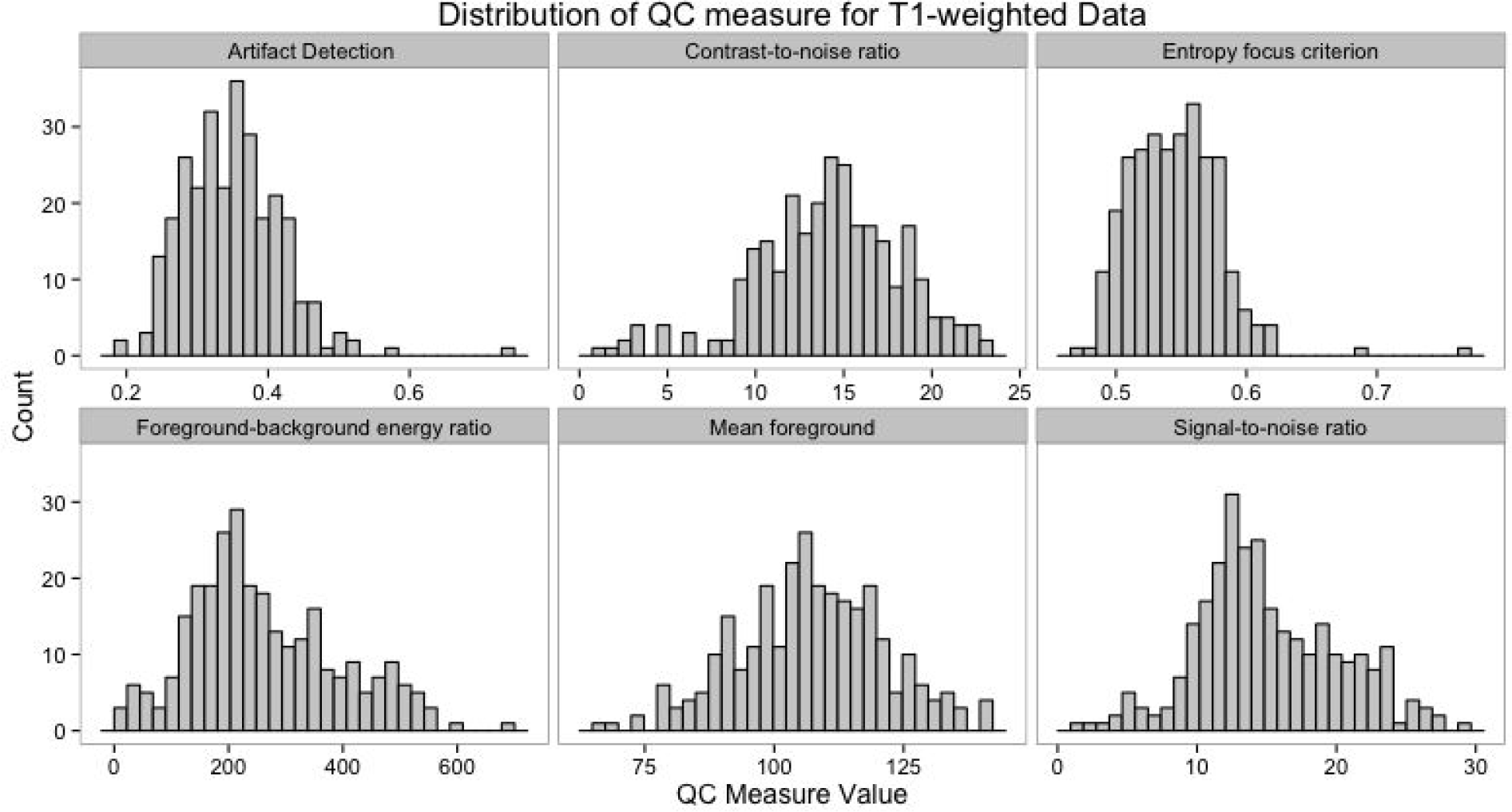
Distribution of selected QC measures for T1w anatomical scans included in dataset.

All Tl-weighted images were skull-stripped ^9^ [AFNI 3dSkullStrip], corrected for intensity inhomogeneity due to B1 variations ^10^ [ANTS N4], and normalized to MNI-152 2 mm template space ^11^ [ANTS]. The background, gray matter, white matter, and cerebrospinal fluid were segmented ^12^ [FSL 5.0.8/FAST], and the resulting segmentations were used to compute the following quality control measures:

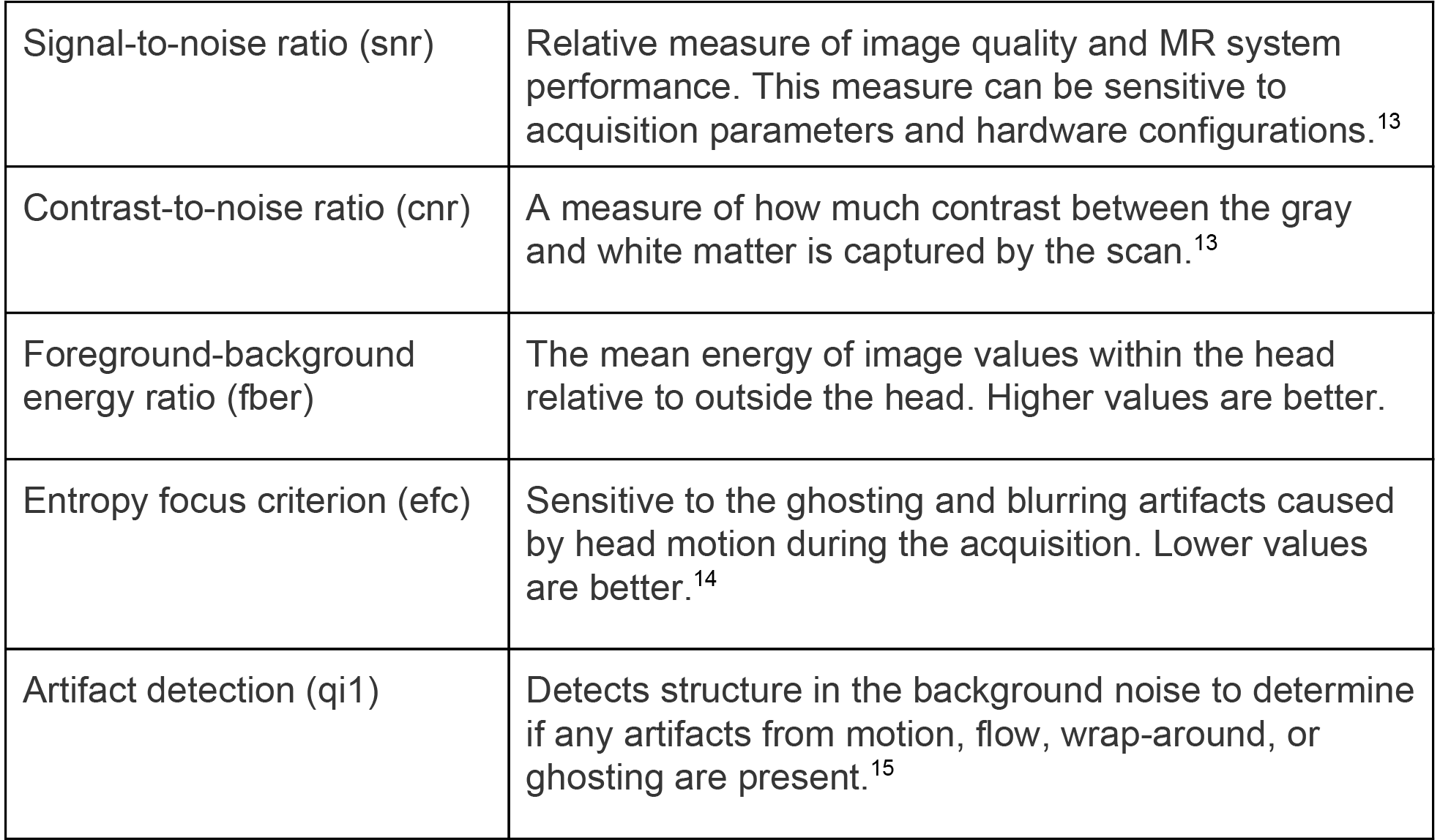

In addition to the quality metrics listed above, summary information about the mean and standard deviation of background, foreground, gray matter, white matter, CSF, and average inhomogeneity bias are provided for comparison across the subject population.

### BOLD/T2*-weighted Functional Scans

**Figure 2:**
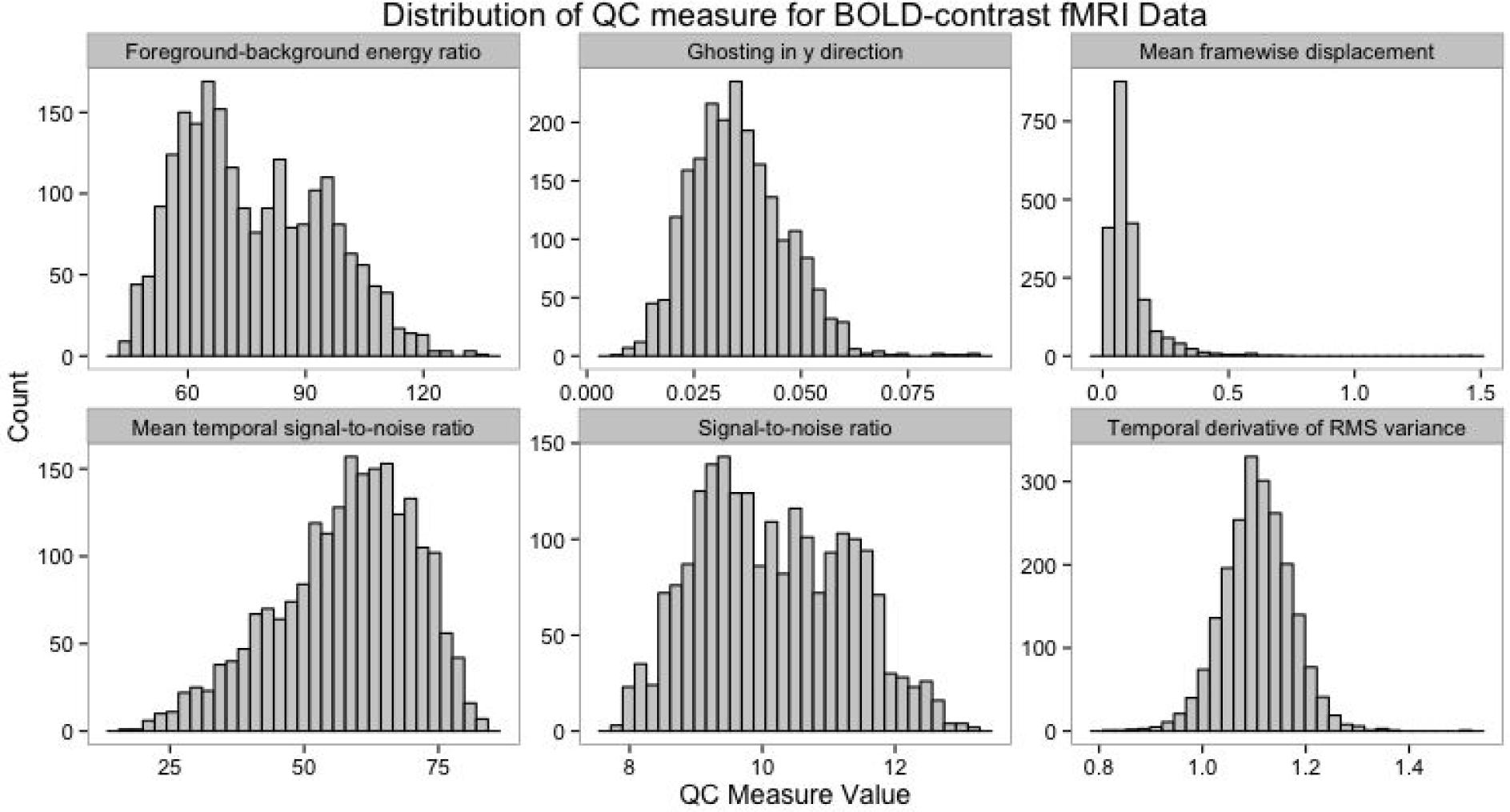
Distribution of selected QC measures for BOLD contrast functional scans included in dataset.

Each BOLD-contrast fMRI scan was corrected for head motion during the acquisition using AFNI ^9^, a brain mask was computed to separate the brain from the skull and outside air ^16^. Then, the following QC metrics were computed:

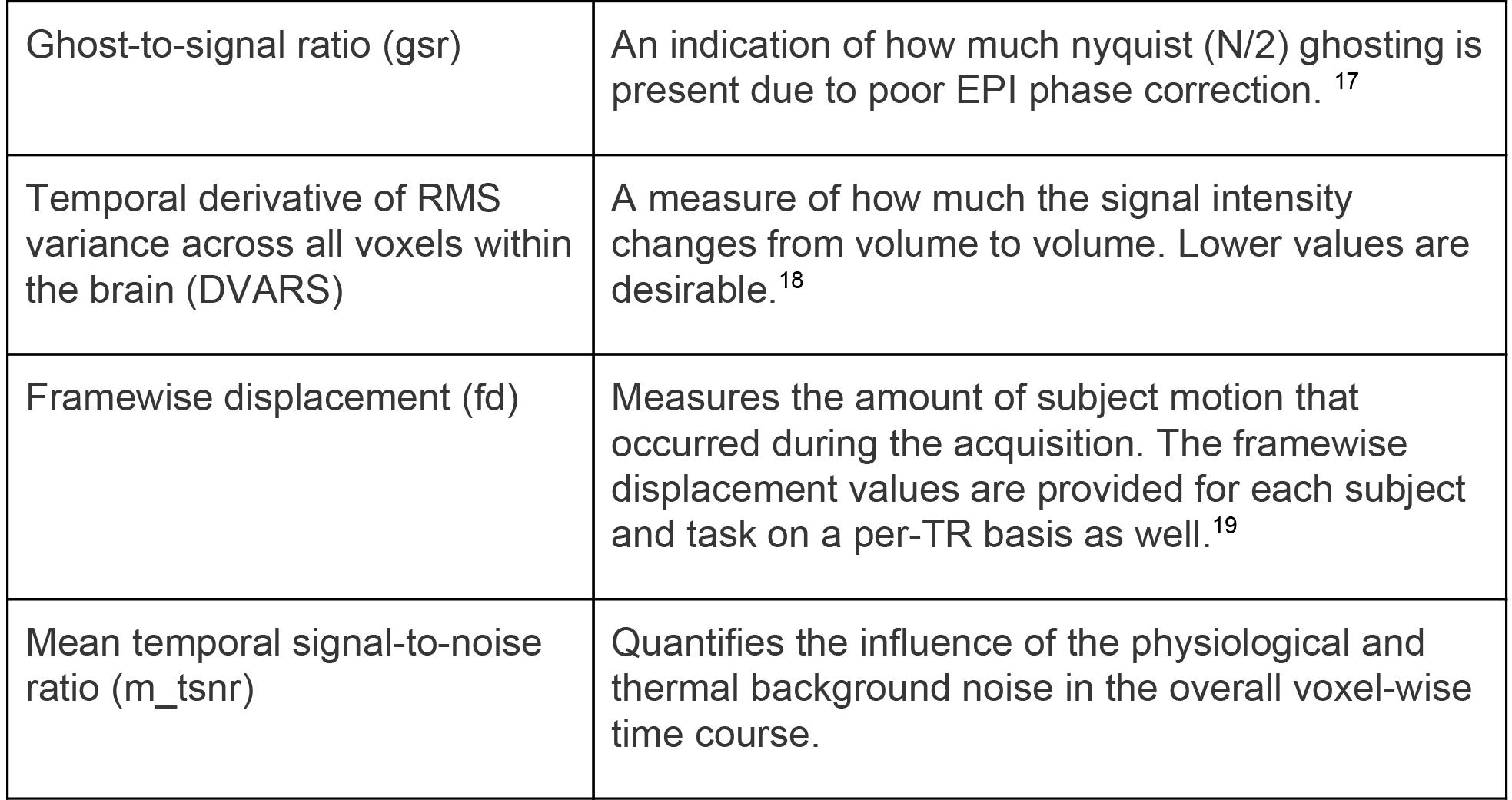

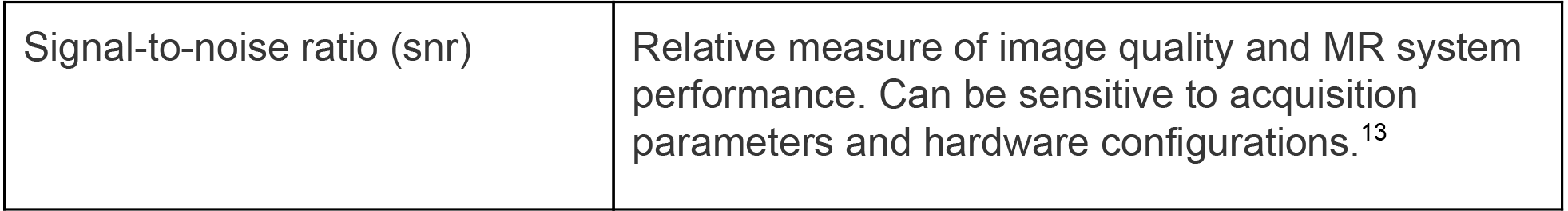

|  |  |
| --- | --- |
| Ghost-to-signal ratio (gsr) | An indication of how much nyquist ($N/2$) ghosting is present due to poor EPI phase correction.<sup>17</sup> |
| Temporal derivative of RMS variance across all voxels within the brain (DVARs) | A measure of how much the signal intensity changes from volume to volume. Lower values are desirable.<sup>18</sup> |
| Framewise displacement (fd) | Measures the amount of subject motion that occurred during the acquisition. The framewise displacement values are provided for each subject and task on a per-TR basis as well.<sup>19</sup> |
| Mean temporal signal-to-noise ratio (m_tsnr) | Quantifies the influence of the physiological and thermal background noise in the overall voxel-wise time course. |
| Signal-to-noise ratio (snr) | Relative measure of image quality and MR system performance. Can be sensitive to acquisition parameters and hardware configurations.<sup>13</sup> |

After the MRIQC was completed, each high-resolution T1-weighted anatomical scan was scrubbed of facial features using the mri_deface program^20^. Visual inspection of all outputs was performed to ensure that the facial features were properly removed to preserve subject privacy.

## fMRI Task Event and Physiological Monitoring Data

fMRI task and event onsets were extracted from the raw matlab log files using R-matlab and are provided as tabular text files. These event files contain the onset, duration, type of trial, reaction time, and task-specific details. The onset time entries in each event file were corrected for the scanner trigger delay to account for extra T1 saturation equalization pre-scans performed by the scanner. In some cases the trigger delay was not explicitly available from the source data. Plots of all event files are provided with the data set for reference.

Similarly, physiological recordings collected during resting state and breath-holding fMRI scans were converted from raw data (Acknowledge format, BioPac) using the Bioread python package (https://github.com/njvack/bioread). The individual recording traces were saved as gzip-compressed tab-separated value files, and the raw data header information (Channel Name, Units, Sampling Rate) was extracted to accompanying JSON files. Plots of physiological traces are provided overlaid with breath-hold instruction timings (ready, hold, rest) for reference.

## Diffusion-weighted Imaging

The diffusion-weighted data were corrected for eddy currents and head motion using the B0 image as the reference, and the motion parameters were saved for later analysis^19^. The corrected volumes were then skull-stripped to remove the background and other non-brain scan regions^16^. Diffusion tensor estimation was performed for each subject, and the mean fractional anisotropy (FA) and average diffusivity (AD) were computed for all brain voxels^21^. Quality assurance was performed using a semi-automated method, including the following steps: confirmed that the b-values and applied directions were the same as expected, calculated mean in-mask FA and average or mean diffusivity (AD, MD), calculated and plotted motion, created a standard deviation image based on regular and motion corrected files, and used regional maps to calculate the percentage of cropped voxels in the occipital lobe, frontal lobe, temporal lobes, cerebellum, and the most superior portion of the brain. Trained individuals then used the results of this script to evaluate scan quality. This included a visual inspection of the FA map, visual inspection of the color map to ensure tracts were coded correctly, a check for cropping in which if more than 10 percent of voxels in a region were cropped, the map was visually inspected to ensure that the cropped region did not encroach on a major tract (i.e., regions that would be included in the FSL DTI skeleton), and the raw data were watched as a movie. Data were flagged for coverage (0=no cropping, 1=minor cropping, 2=severe unuseable cropping), motion (based on watching raw data as movie, and on motion plots), tensor direction flags (based on b-value and direction calculations, and on observation of the color map), artifact flags (including noise, striping, and the vibration artifact that affected many Siemens Trio systems during this time period). An overall quality score from 1–4 was generated from these measures, in which 1=excellent, 2=good, 3=fair (both 2 and 3 may be useable, but depending on analysis may want to consider the reason for the decreased score), and 4=unuseable (all individuals with vibration artifacts are in this category, along with other irreconcilable problems), 5=not evaluated. Ratings are included with the data download.

**Figure 3:**
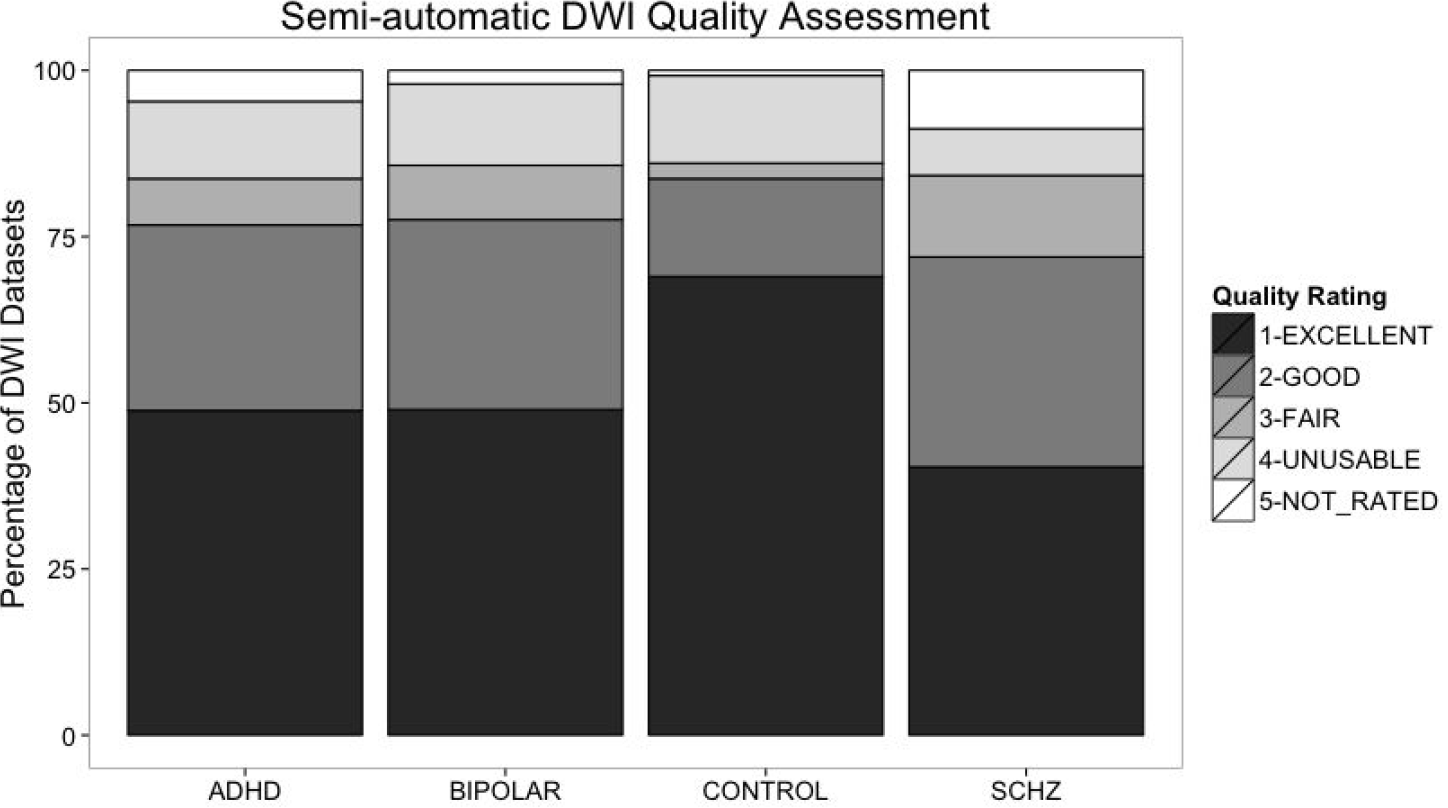
Results from the semi-automated DWI quality assessment, by subject group.

## Usage Notes

All data are made available under the Public Domain Dedication and License v1.0 (http://www.opendatacommons.org/licenses/pddl71.0/), which places no restrictions on the usage of the data. We expect that users of the data will follow the ODC Attribution-Sharealike Community Norms (http://opendatacommons.org/norms/odc-by-sa/), which state that any derivative works should be shared via an equally open license, and that the creators of this dataset should be credited by citation of the present data descriptor.

We ask that users with questions please use the NeuroStars Forum (https://neurostars.org) and attach the tags "openfmri" and "ds000030" in order to discuss and comment on this dataset.

## Data Citations

R. A. Poldrack et al., *UCLA Consortium for Neuropsychiatric Phenomics LA5c Study*. https://openfmri.org/dataset/ds000030/

## Acknowledgments

This work was supported by the Consortium for Neuropsychiatric Phenomics (NIH Roadmap for Medical Research grants UL1-DE019580, RL1MH083268, RL1MH083269, RL1DA024853, RL1MH083270, RL1LM009833, PL1MH083271, and PL1NS062410). Preparation of the data in the BIDS format was supported by a grant from the Laura and John Arnold Foundation. Thanks to many research assistants who helped with data collection: especially Angelica Bato and Eric Miller who were primarily charged with scanning procedures, and others who participated in recruitment, clinical assessment, cognitive assessment, translation, task programming, and assisting with scanning, including Hannah Al-Sodani; Oren Boxer; Xavier Cagigas; Rachel Casas; Alan Chang; Jennifer Erickson; Winifred Flach; Christina Fong; Chelsea Gilbert; Samantha Hemingway; David Kaufman; Milky Kohno; Rachel Lavian; Evan Lutkenhoff; Carla Orieta; Katy Preciado; Cesia Toledo; Shahaf Tuler; Jessica Valluzzi; Jonathan Yang, Anna Xu, Amira Ibrahim, Aron Jacobsen, Nathalie De Shetler, Borja Izaguirre, Tyler Morgan, and Joey Contreras

## Author contributions

RAP: conception and design of study and data sharing plan, drafting of the article. EC: conception and design of study, acquisition and analysis of data, critical review and final approval of the version submitted. WT: curation, critical review and final approval of the version submitted. KJG: design of data sharing plan, critical review and final approval of the version submitted. KHK: conception and design of study, acquisition and analysis of data, critical review and final approval of the version submitted. JAM: conception and design of study, final approval of the version submitted. FWS: conception and design of study, final approval of the version submitted. NBF: conception and design of study, final approval of the version submitted. TDC: conception and design of study, final approval of the version submitted. RMB: conception and design of study and data sharing plan, critical review and final approval of the version submitted.

---

**Table S1.**
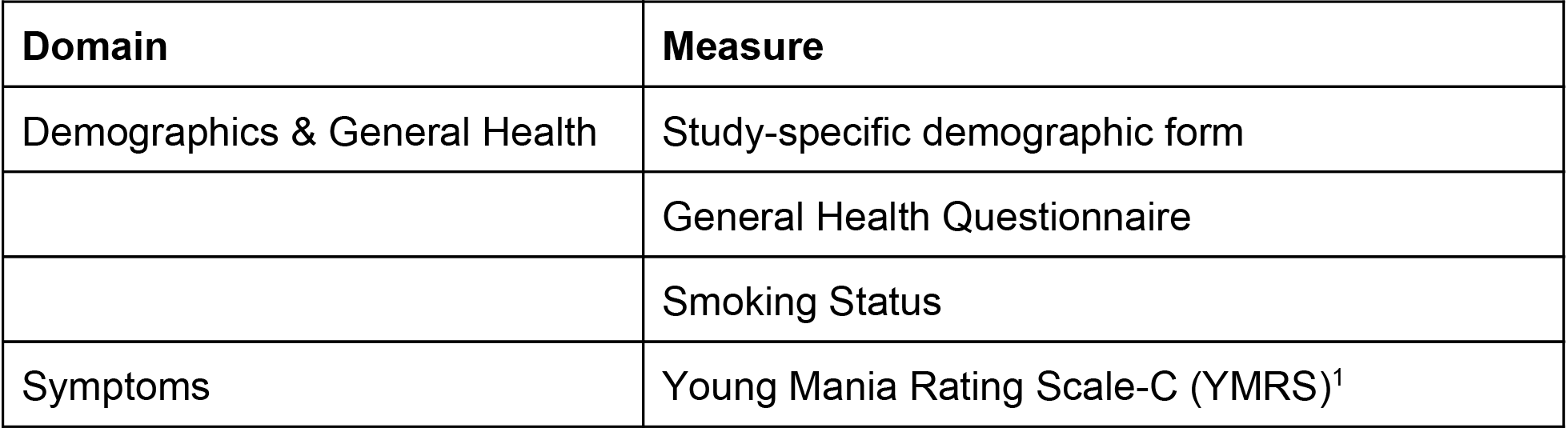
Behavioral session assessments.

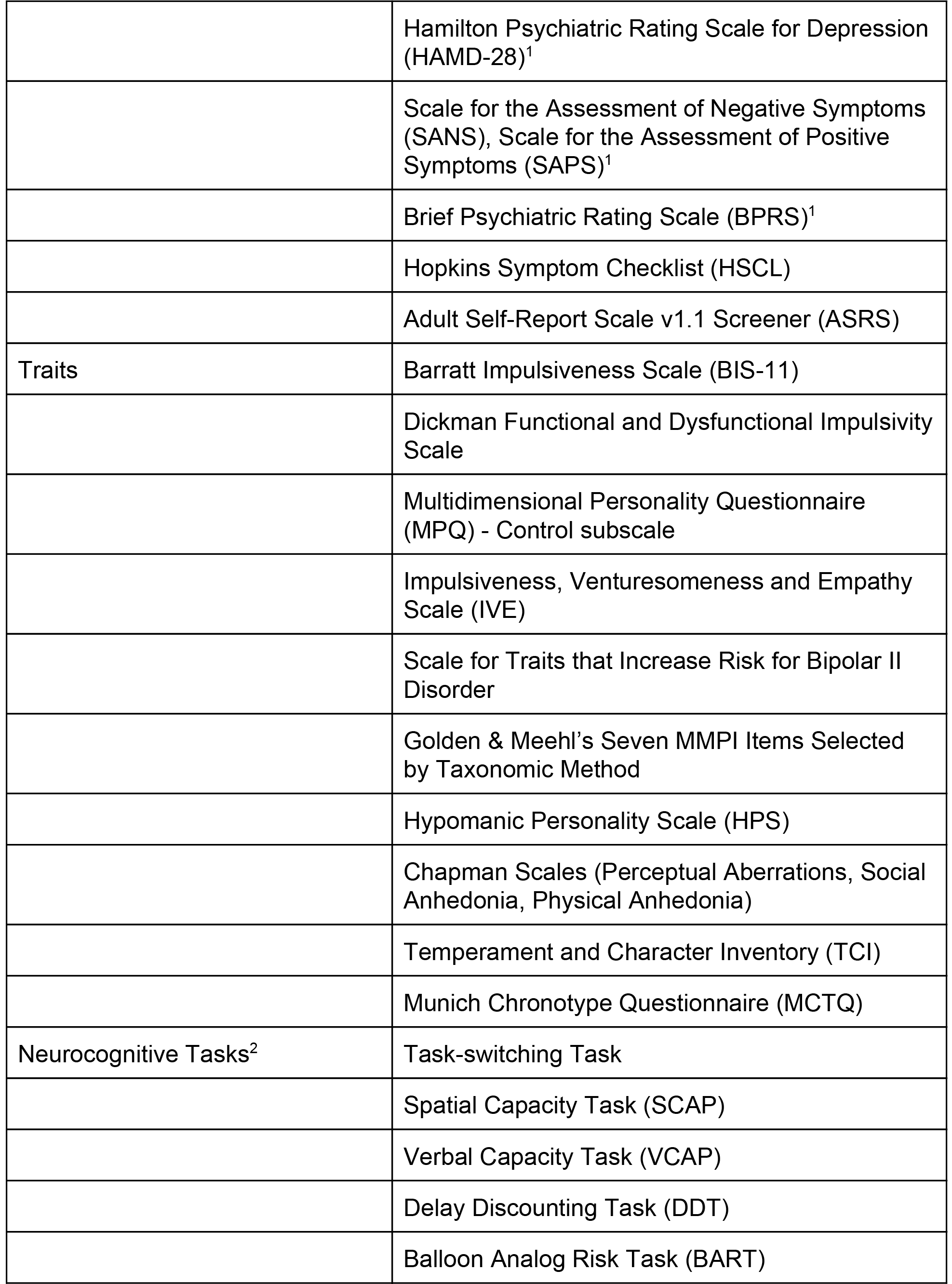

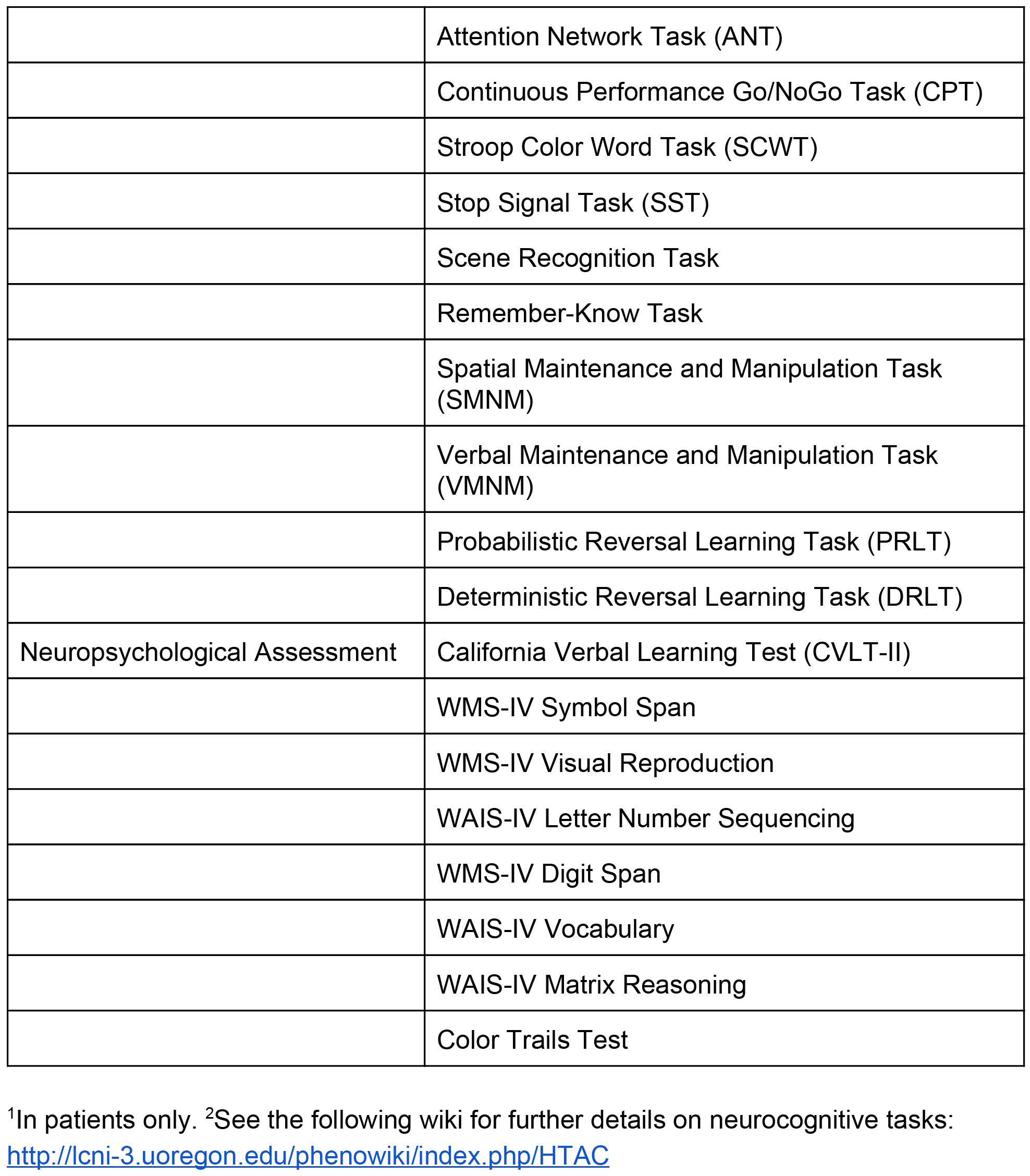

